# Evolution of Secondary Cell Number and Position In the Drosophila Accessory Gland

**DOI:** 10.1101/2022.11.25.517994

**Authors:** Yoko A. Takashima, Alex C. Majane, David J. Begun

**Author notes:** Corresponding Author(s) (YAT), (DJB).

## Abstract

In animals with internal fertilization, males transfer gametes and seminal fluid during copulation, both of which are required for successful reproduction. In *Drosophila* and other insects, seminal fluid is produced in the paired accessory gland (AG), the ejaculatory duct, and the ejaculatory bulb. The *D. melanogaster* AG has emerged as an important model system for this component of male reproductive biology. Seminal fluid proteins produced in the *Drosophila* AG are required for proper storage and use of sperm by the females, and are also critical for establishing and maintaining a suite of short- and long-term postcopulatory female physiological responses that promote reproductive success. The *Drosophila* AG is composed of two main cell types. The majority of AG cells, which are referred to as main cells, are responsible for production of many seminal fluid proteins. A minority of cells, about 4%, are referred to as secondary cells. These cells, which are restricted to the distal tip of the *D. melanogaster* AG, may play an especially important role in the maintenance of the long-term female post-mating response. Many studies of *Drosophila* AG evolution have suggested that the proteins produced in the gland evolve quickly, as does the transcriptome. Here, we investigate the evolution of secondary cell number and position in the AG in a collection of eight species spanning the entire history of the *Drosophila* genus. We document a heretofore underappreciated rapid evolutionary rate for both number and position of these specialized AG cells, raising many interesting questions about the developmental, functional, and evolutionary significance of this variation.

## Introduction

Along with sperm, *Drosophila* males transfer seminal fluid proteins (often referred to as Acps or Sfps) to females. Transfer of molecules from male to female provides a fascinating example of male-female co-operation and conflict (reviewed in Wolfner 2011). Female reproduction requires seminal fluid, yet this requirement exposes females to male adaptations that may subvert their reproductive interests (Rice 1996, Holland and Rice 1999, Lung and Wolfner 1999, Civetta and Clark 2000, Fiumera *et al*. 2006, Sirot *et al*. 2015). Seminal fluid may also mediate competitive interactions between the sperm of multiple males in the female reproductive tract (Harshman and Prout 1994, Clark *et al*. 1995, Fiumera *et al*. 2007). Significant polymorphism of genes affecting seminal fluid function is evident in the rapid laboratory evolution of male toxicity toward females and the rapid evolution of male sperm displacement phenotypes in experimental populations of *Drosophila melanogaster* (Rice 1996, Holland and Rice 1999, Hollis *et al*. 2019). The outcome of sperm competition is also influenced by female genetic variation and the interaction of genetic variation in the two sexes (Clark and Begun 1998, Clark *et al*. 1999, Giardina *et al*. 2011).

*Drosophila* Sfps are produced in the accessory gland (AG), ejaculatory duct, and ejaculatory bulb. The molecular products of the seminal fluid producing organs exhibit rapid evolution, as expected given their role in sexual conflict (Swanson and Vacquier 2002). Sfp protein sequences often evolve very quickly (Coulthart and Singh 1988), sometimes under the influence of directional selection (Begun *et al*. 2000; Swanson et al. 2001; Wagstaff and Begun 2005a; Haerty et al. 2007). Expression levels of AG-biased genes may vary widely even between closely related species (Begun and Lindfors 2005, Ahmed-Braima *et al*. 2017, Garlovsky et al. 2020), and the AG transcriptome exhibits rapid turnover due to gene presence/absence variation (Wagstaff and 2005b, Mueller *et al*. 2005; Begun *et al*. 2006) or the expression of different homologues in the AG of different species (Ahmed-Braima et al. 2017; Cridland *et al*. 2020, 2022; Hurtado *et al*. 2022).

The AG is composed of two major cell types. In *D. melanogaster*, main cells constitute about 96% of the cells, which are the primary producers of the secreted proteins required for fertility, and induce both short- and long-term changes to female post-copulatory physiology and behavior (Manning 1967, Kalb *et al*. 1993, Harshman and Prout 1994, Xue and Noll 2000, Neubaum and Wolfner 1999, Ravi Ram and Wolfner 2007, 2009). The secondary cells (estimated to be 4% of the cells) are clustered in the distal portion of the organ (reviewed in Wilson *et al*. 2017). Their secreted products appear to play an important role in maintaining female post-copulatory responses (Sitnick *et al*. 2016, Hopkins *et al*. 2019, reviewed Wilson *et al*. 2017). Secondary cells (SC) play a role in the secretion of extracellular vesicles (*i*.*e*., exosomes), which bind to sperm, interact with the female reproductive tract epithelium, and play a role in the long-term mating response of females (Corrigan *et al*. 2014). Recent RNA-seq analysis of the SC has revealed that its transcriptome is distinct from that of the main cells and also evolves in a distinct manner (Immarigeon *et al*. 2021, Majane *et al*. 2022). Given that male reproductive proteins and transcriptomes evolve quickly and that mating systems vary dramatically across *Drosophila* (Markow 1996), whether most basic conclusions about the AG as described in the *D. melanogaster* model apply generally to *Drosophila* is an open question. A report by Taniguchi *et al*. (2012), which focused on AG main cell binucleation in multiple *Drosophila* species, provided incidental data (their Fig. 1) strongly suggesting that Drosophila secondary cell number evolves, perhaps quickly. The work reported here quantitatively investigates this possibility by systematically estimating secondary cell number in a collection of *Drosophila* species.

**Fig 1:**
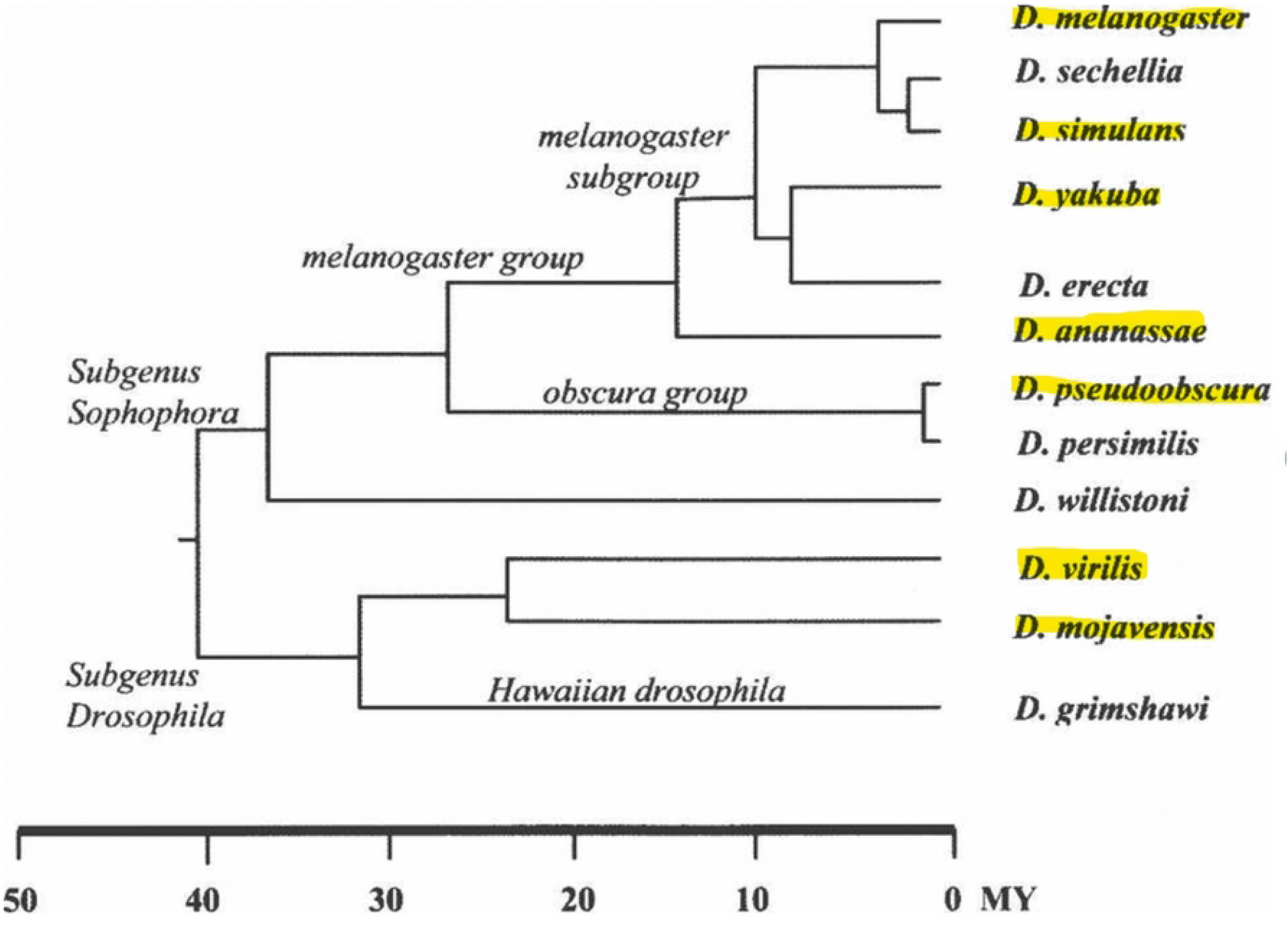
Phylogenetic Tree of *Drosophila* Species (Powell 1997)

## Materials and Methods

Fly Stocks: *D. melanogaster* RAL 517, RAL 399, RAL 360 (Mackay *et al*. 2012), *D. yakuba* Tai18E2 (Begun *et al*. 2007), *D. simulans w*^*501*^ (Begun *et al*. 2007), *Drosophila hydei, D. virilis* (gift of S. Lott), *D. mojavensis* (Drosophila Species Stock Center #s 0218.15 and 0218.17), *D. ananassae* 11-4 (gift of Brandon Cooper, University of Montana), *D. pseudoobscura* SLC 10 (gift of S. Schaeffer, Pennsylvania State University). All stocks were maintained in a 25C incubator on a 12:12 hr light:dark cycle.

The phylogenetic relationships of these species is depicted in FIG. 1 (FIGURE 1 - Phylogeny of *Drosophila* species used). Four of these species (*D. melanogaster, D. simulans, D. yakuba*, and *D. ananassae*) are from the *melanogaster* group; two (*D. hydei* and *D. mojavensis*) are from the *repleta* group; one (*D. pseudoobscura*) is from the *obscura* group, and one (*D. virilis*) is from the *virilis* group. The species were sampled to provide information about shorter and longer time scale evolutionary change. On a long timescale, the most recent common ancestor of all eight species existed about 40-50 million years ago (Powell 1997, Tamura *et al*. 2004). On a shorter timescale, divergence between *D. melanogaster* and *D. simulans* is about 2-3 million years (Obbard *et al*. 2012). Three inbred lines of *D. melanogaster* were used to investigate the possibility of genetic variation affecting SC number in this species.

### Dissecting, Fixing, and Staining Glands

Virgin males were collected and then dissected two days after their estimated age of sexual maturity, which varies among species (Markow 1996). *D. melanogaster, D. simulans*, and *D. yakuba* were dissected two days post-eclosion. *D. hydei* were dissected 10 days post eclosion. *D. mojavensis, D. pseudoobscura, D. ananassae*, and *D. virilis* were dissected eight days post eclosion.

Dissected accessory glands were fixed in 4% paraformaldehyde for 30 minutes at room temperature, and then washed with 1X PBS. The glands were treated with a 1:100 dilution of 10mg/ml RNAse in PBT for 30 minutes at room temperature, and stained using propidium iodide at a concentration of 1:1000 for 30 minutes at room temperature (Taniguchi et al., 2012). Glands were then mounted on glass slides with spacers - Scotch tape layered and cut into squares that would mark where the corners of the 0.5mm coverslip would lie. Spacers were necessary to prevent compression of the gland. One gland of the two was imaged using a Leica confocal microscope at 20X (dry) and 40X (oil) magnification with 75-100 z-stacks per gland. The Z step size was approximately 1.8-2.0μm. Images were stitched together using the Pairwise Stitching of Images (Preibisch et al. 2009) or were merged at the Leica confocal upon image acquisition using the tilescan function with 10% overlap.

### Secondary Cell Quantification

Due to the nature of the propidium iodide staining and the structure of the cell itself, SCs were easily distinguished from main cells by their characteristic large, dark vacuoles (FIG. 2). SCs were counted manually across z-stack images using the Cell Counter Plugin in FIJI (De Vos et al 2001). After initializing a .tif file in this plugin, the “show all” function allowed us to track and count each SC throughout the z plane. Two arbitrary counters, which were conveniently different colors, were used to tally the different secondary cells to properly ensure that double counting did not occur. One counter tracked one side of the epithelial layer, and after going through the lumen of the gland; the second counter was used to track cells on the other epithelial layer. The plugin then tallied the number of times the counter was used. Totals from the two counters were summed to estimate the number of secondary cells in a gland. Distinguishing between each epithelial layer was possible due to the spacers on the slide to prevent the compression of the tissue.

**Fig 2.**
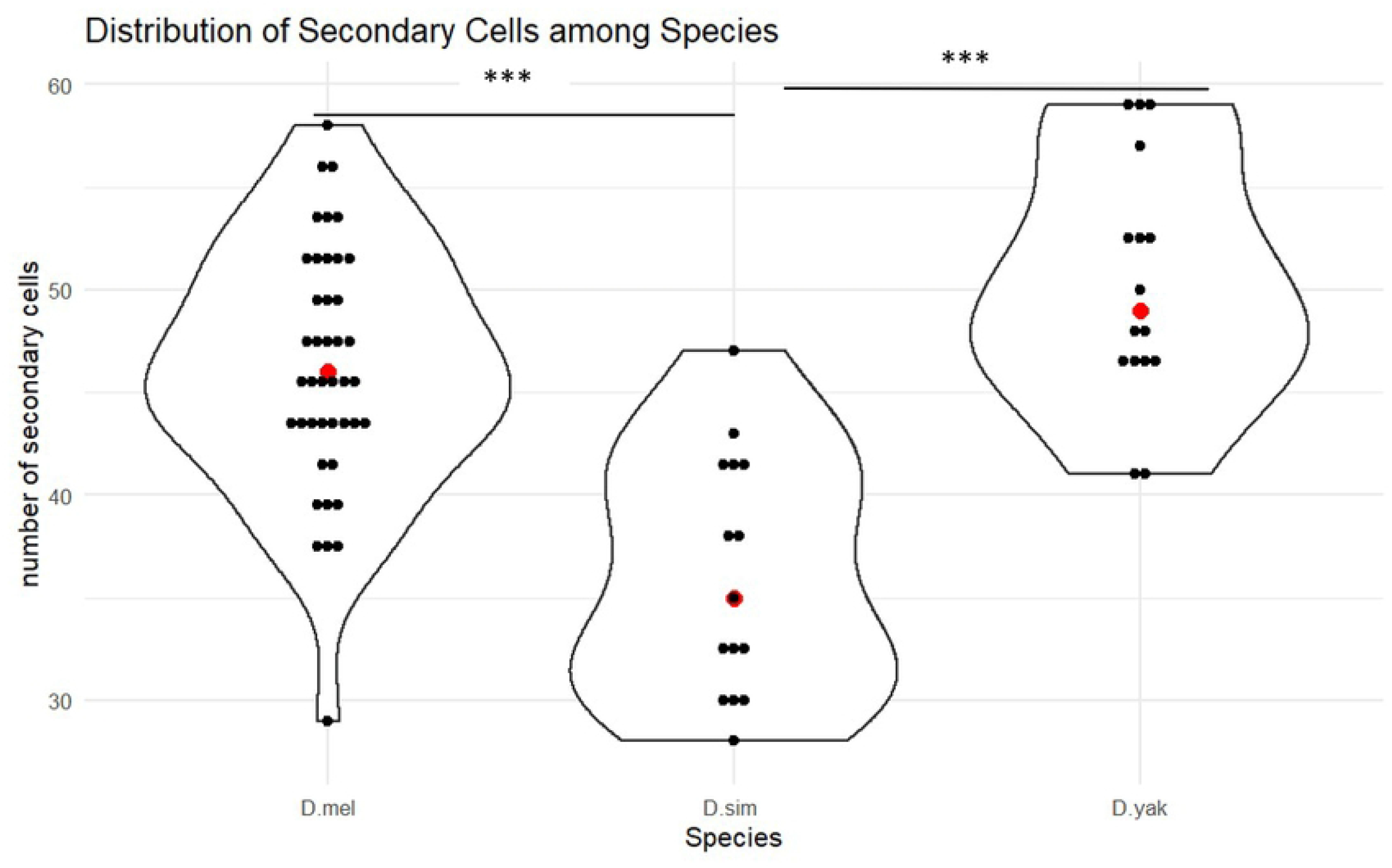
Distribution of Secondary Cells Across the Melanogaster subgroup. Violin plot showing distributions of number of secondary cells in each species. Red dot = mean. Asterisks indicate p-values from ANOVA. * < 0.05, ** < 0.01, *** < 0.001

### Accessory Gland Length, Area, and Volume Measurements

AG gland measurements were done using FIJI. Gland length measurements were done by using the freehand tool and drawing a line directly down the middle of the gland spanning from the top of the ejaculatory duct and the tip of the gland. The measurements were taken in pixels following Bangham *et al*. (2002) and Baker *et al*. (2003).

AG area was taken by using the lasso tool on FIJI and drawing a perimeter around the glands. Once a perimeter of the gland was drawn, measurements were taken and the area in microns was generated following Bangham *et al*. (2002) and Baker *et al*. (2003).

### Statistics

Statistics were done using R/Rstudio version 2022.02.3 Build 492. “Prairie Trillium” Release (1db809b8, 2022-05-20) for Windows. For ANOVA and Tukey’s Honest Significant Difference Test the package multicomp was used (Hothorn 2008). Means and standard deviations were done using the statistical packages in R (R Core Team 2020)

## Results and Discussion

Table 1 presents our estimates of SC number in eight species. Previous estimates of SC number in *D. melanogaster* were roughly 43 SC per gland (Bertram *et al*. 1992). Our results provide a similar, though slightly greater estimate of SC number per gland, and also provide a quantitative measure of variation among genotypes. The three *D. melanogaster* inbred lines we assayed, which originated in Raleigh, North Carolina (Mackay *et al*. 2012), were not significantly heterogeneous for SC number. Thus, while there may be genetic variation for SC number currently segregating in *D. melanogaster*, we observed no evidence for this in our small sample. It is worth noting that a range of SC numbers were observed within each inbred RAL genotype (e.g., for RAL 517 the range across individuals was 29 to 56). While some of this variation could represent measurement error, developmental variation in the process leading to SC differentiation could also contribute. *D. simulans*, the sister species to *D. melanogaster*, had significantly fewer SC (FIG 2). This provides strong evidence that SC-number can evolve substantially, even on short timescales. Of course, the observation of species differences provides *primae facie* evidence for genetic variation affecting SC number. While our estimate of *D. yakuba* SC number is greater than either *D. melanogaster* or *D. simulans*, it is not significantly different from *D. melanogaster*. Thus, the most parsimonious explanation for the observed variation in SC number in the *melanogaster* subgroup is a recent decrease in SC number in *D. simulans*. All samples from the three *melanogaster* subgroup species exhibited the well documented spatial patterning previously observed in *D. melanogaster*, with all SC distributed in the distal tip of the gland (FIG. 3).

**Table 1:** Secondary Cell Number for Eight *Drosophila* Species.

| Species (# glands) | Mean # of SC<br>(std.dev) |
| --- | --- |
| *D. melanogaster* (n=42) | 46 (6) |
| *D. simulans* (n=15) | 36 (5) |
| *D. yakuba* (n=16) | 50 (5) |
| *D. ananassae* (n=15) | 39 (7) |
| *D. pseudoobscura* (n=7) | 23 (6) |
| *D. mojavensis* (n=15) | 239 (44) |
| *D. virilis* (n=13) | 122 (20) |
| *D. hydei* (n=15) | 259 (54) |

**Fig 3.**
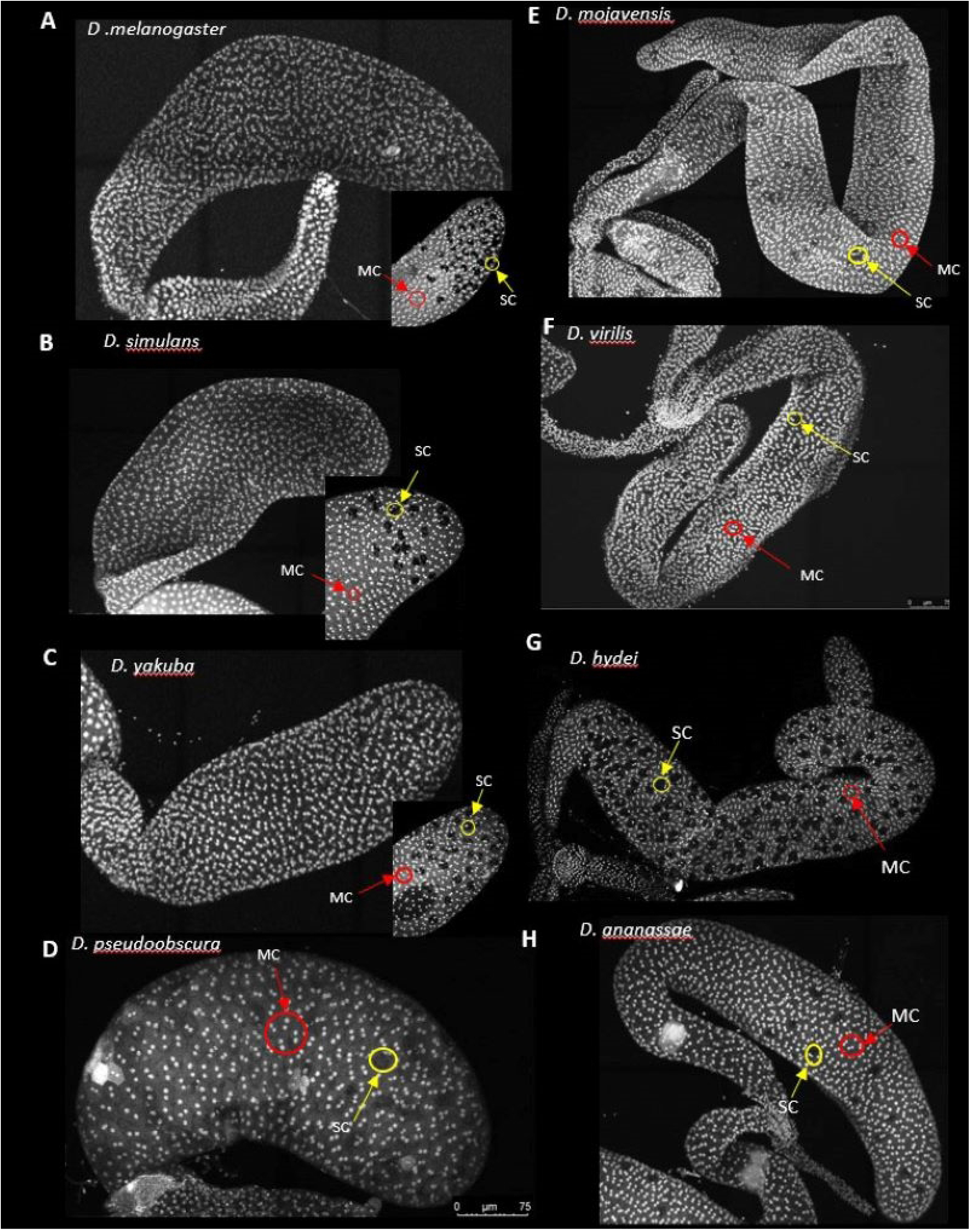
Confocal Images of Representative AGs For Eight *Drosophila* Species. (A-C) Accessory glands imaged at 40X, tilescan with 10% overlap, max projection confocal images of *D. melanogaster, D. simulans, D. yakuba*. The inset is a 40X image of the distal tip of those species. (D-H) are 40X tilescan images, 10% overlap, max projection confocal images of D) *D. pseudoobscura*, E) *D. mojavensis*, F) *D. virilis*, G*) D. hydei*, H) *D. ananassae*

*D. ananassae* is the nearest outgroup for the *melanogaster* subgroup in our sample. *D. ananassae* mean SC number was significantly lower than the mean for the *melanogaster* subgroup (FIG 2), but without formal phylogenetic modeling of the trait we cannot infer the ancestral SC number of the *melanogaster* group and thus the direction of evolution in the branch leading to *D. ananassae* and the branch leading to the *melanogaster* subgroup. Despite the similar number of SC cells in *D. ananassae* and the *melanogaster* subgroup, there is a clear difference in their spatial distribution in the gland. *D. ananassae* differs from the three *melanogaster* subgroup species in that the SC are not restricted to the distal tip of the gland. Instead they appear to be roughly homogeneously distributed throughout the distal half of the gland (FIG 3).

*D. pseudoobscura*, our only representative of the *obscura* group, is sister to the *melanogaster* group. It exhibits fewer SC than any *melanogaster* subgroup species, though the difference is not significant. Much like the *melanogaster* subgroup, the secondary cells are enriched in the distal tip of the gland. The morphology of the *D. pseudoobscura* accessory gland was clearly shorter and about the same width (mean length and width, 546 microns and 191 microns, respectively) compared to the *melanogaster* subgroup (mean length and width, 1426 microns and 189 microns, respectively).

*D. hydei* and *D. mojavensis*, sister species in our sample, belong to the *repleta* group, a diverse clade of primarily cactophilic flies that originated about 20-30 MYA, most likely in South America (Patterson and Stone 1952, Oliveira *et al*. 2012). The reproductive biology of these flies differs from several other *Drosophila* species in multiple ways. For example, *D. mojavensis* and *D. hydei* share high re-mating rates compared to most other *Drosophila* species, with *D. hydei* apparently having a particularly high rate (reviewed in Markow 1996). *D. hydei* also has very long sperm and testis compared to most flies (Pitnick and Markow 1994). We observed that these two species have extraordinarily high numbers of SC per gland, roughly six times as many as *melanogaster* (Table 1). Given that the *D. hydei* gland is only about four times the size of *D. melanogaster* (Table 2), it appears that the species differences in SC number cannot be explained solely by gland size variation, which is driven primarily by gland length differences.

**Table 2:** Mean Area and Length of AGs (microns)

| Species (# glands) | Area in microns (std.dev) | Length in microns (std.dev) |
| --- | --- | --- |
| *D. melanogaster* (n=9) | 129476 (24130) | 801(90) |
| *D. hydei* (n=7) | 514505 (633262) | 2017(854) |

Supporting this conclusion, in striking contrast to the *melanogaster* group, *repleta* group flies exhibit SC distributed homogeneously throughout the gland (FIG 3). Interestingly, the *repleta* group is the only clade of *Drosophila* thought to be lacking the sex-peptide gene (McGeary and Clearly 2020, Hopkins and Perry 2022), which codes for a protein critical for proper sperm storage and use in *D. melanogaster* (Chen *et al*. 1988). Whether the absence of sex-peptide, the increase in number of SC, and the homogeneous physical distribution of SC in the AG in the *repleta* group are functionally related is an interesting question.

*D. virilis*, which is sister to the *repleta* group (FIG. 1), also exhibits substantially greater numbers of SC than the *Sophophora* species, though many fewer than *repleta* group flies (FIG 4). Thus, the sampled taxa are suggestive of greater numbers of SC per gland for the *Drosophila* group than for the *Sophophora* group (FIG. 1), though additional sampling would be required to be confident of this inference. Similar to the *repleta* group species, *D. virilis* SC appear to be distributed homogeneously throughout the gland (FIG. 3). In general, our data support the idea that restriction of the SC to the distal tip of the accessory gland in the *D. melanogaster* model system is a highly derived trait and not characteristic of most *Drosophila* lineages. The developmental basis of SC number and position, as well as their functional consequences and evolutionary processes driving divergence of these traits, are unknown.

**Fig 4.**
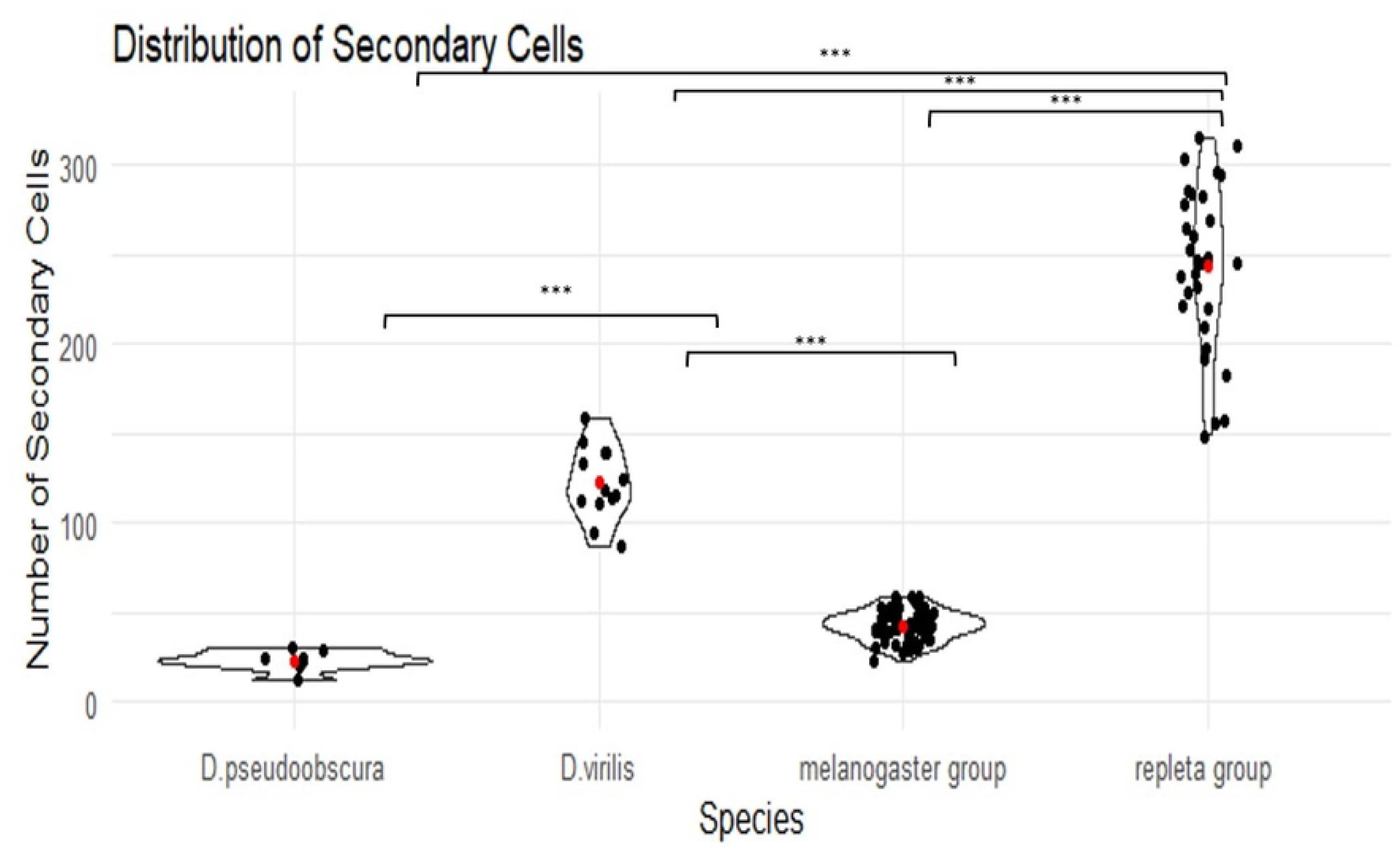
Distribution of Secondary Cells Across Species. Violin plot showing the distribution of secondary cells among the melanogaster, repleta, virilis, and pseudoobscura groups. Red dots indicate the mean number of secondary cells. Asterisks indicate significance p-values from ANOVA: * = 0.05, ** = 0.01, *** = 0.001

## Conclusion

While formal phylogenetic analysis of SC number and position in the accessory gland would be necessary to make strong, specific inferences on rates of phenotypic evolution of SC number on particular branches of the *Drosophila* phylogeny, the results reported here leave no doubt that SC number evolves quickly in the genus. Additionally, the physical distribution of SC in the gland also evolves, as most species do not appear to share the stereotypical clustering of SC in the tip of the gland seen in the *D. melanogaster* model. Further investigation of the patterns presented here promise to reveal new functional and evolutionary attributes of accessory gland diversification and its broader connections to mating system biology in the genus.

## Acknowledgements

This work was supported by the National Institutes of Health grant NIGMS R35 GM134930 to DJB and a National Science Foundation Graduate Research Fellowship to ACM. We thank D. Sarikaya for microscopy and image analysis advice and feedback. We thank Noelia Carrasquila and Maria Florencia Ercoli for assistance with the confocal microscope and the UC Davis Department of Plant Pathology for access to it. We thank Kiichiro Taniguchi for sharing his Propidium Iodide staining protocol.

